# No priming, just fighting – endophytic yeast attenuates the defense response and the stress induced by Dutch elm disease in *Ulmus minor*

**DOI:** 10.1101/2022.01.29.478285

**Authors:** J. Sobrino-Plata, C. Martínez-Arias, S. Ormeño-Moncalvillo, I. Fernández, C. Collada, L. Gil, C.M.J. Pieterse, J.A. Martín

## Abstract

One century after the first report of Dutch Elm Disease (DED), there is still no practical solution for this problem threatening European and American elms (*Ulmus* spp.). The long breeding cycles needed to select resistant genotypes and the lack of efficient treatments keep disease incidence at high levels. In the present work, the expression of defense-related genes to the causal agent of DED, *Ophiostoma novo-ulmi*, were analyzed in in vitro clonal plantlets from two DED-resistant and two DED-susceptible *U. minor* trees. In addition, the effect of the inoculation of an endophytic pink-pigmented yeast (*Cystobasidium* sp.) on the plant’s defense system was tested both individually and in combination with *O. novo-ulmi*. The multifactorial nature of the resistance to DED was confirmed, as no common molecular response was found in the two resistant genotypes. However, the in vitro experimental system allowed to discriminate the susceptible from the resistant genotypes, showing higher levels of oxidative stress and phenolic compounds in the susceptible genotypes after pathogen inoculation. Inoculation of the endophyte before *O. novo-ulmi* attenuated the plant molecular response induced by the pathogen and moderated oxidative stress levels. Niche competition, endophyte-pathogen antagonism, and molecular crosstalk between the host and the endophyte are discussed as possible mechanisms of stress reduction. In sum, our results confirm the multifactorial nature of DED resistance mechanisms and highlight the possibility of using certain endophytic yeasts as biological tools to improve tree resilience against biotic stress.

## Introduction

One hundred years ago, the Dutch phytopathologist Bea Schwarz, led by Professor Johanna Westerdijk (Boonekamp et al. 2019), described for the first time the Dutch Elm Disease (DED) (Schwarz 1922). Since then, European elm populations have suffered a massive reduction due to the constant threat of this vascular wilt disease caused by the ascomycete fungi *Ophiostoma ulmi* and *O. novo-ulmi*. Bark beetles are responsible for transmitting fungal spores into healthy elm trees, where they germinate and spread into the xylem vessels inducing their blockage and embolism. Thus, water transport is critically hindered resulting in foliage wilting and tree death (Brasier 1991). Location, characterization, and propagation of pure *Ulmus minor* germplasm have become a priority to conserve and restore lost elm populations (Martín et al. 2019b). In Spain, large efforts have been invested to breed resistant *U. minor* genotypes by screening plant materials from all along the country. By now, seven *U. minor* genotypes have been registered as resistant base materials for forest use (Martín et al. 2015). These resistant genotypes are also valuable materials for elucidating the basis of *U. minor* resistance to DED (Li et al. 2016; Pita et al. 2018).

Chemical and anatomical factors partially explain DED resistance. For instance, the accumulation of lignin, suberin and mansonones in response to *O. novo-ulmi* infection is usually higher in DED-resistant than in DED-susceptible genotypes (Duchesne et al. 1986; Martin et al. 2005; Martín et al. 2008), as well as the constitutive proportion of cellulose and hemicellulose in xylem tissues (Li et al. 2016). In contrast, DED-susceptible genotypes tend to possess wider xylem vessels and a higher proportion of large vessels than resistant ones, enabling fungal dispersal throughout the plant (Solla and Gil 2002; Martín et al. 2013a), although recent research has shown that some resistant genotypes also form wide vessels (Martín et al. 2020). Beyond these differences, other key resistance traits are possibly encrypted in the genetic code of each genotype. The genetic basis of the *U. minor* response to *O. novo-ulmi* has just started to be elucidated by using classical and novel “omic” technics, such as 454-sequencing (Perdiguero et al. 2015) and RNA-sequencing (authors, unpublished results). A recent work by Perdiguero et al. (2018) described the molecular responses activated over time upon *O. novo-ulmi* infection in a highly DED-susceptible clone. The results pointed to defense mechanisms that are regulated by the salicylic acid (SA) pathway. SA-dependent defense responses are typically triggered by biotrophic pathogens (Pieterse et al. 2009). *O. novo-ulmi* is considered a hemi-biotrophic pathogen, because it has an initial biotrophic phase during the vascular colonization of the xylem, followed by a necrotrophic phase during more advanced disease stages (Martín et al. 2012; Sherif et al. 2017). SA-dependent defenses can be activated systemically to distal parts of the plant through molecules such as methyl-SA (MeSA) or glycerol-3-phosphate (G3P), where they play a role in the activation of systemic acquired resistance (SAR) (Fu and Dong 2013). Both, local and systemic defense activation is associated with the accumulation of pathogenesis related (PR) proteins, some of which possess antimicrobial activities against a broad range of pathogens (Bari and Jones 2009).

Plant-microbe symbiotic associations provide plants with higher phenotypic plasticity to changing environments, including biotic and abiotic stresses (Liu et al. 2020). Enhancement of nutrient acquisition and defensive metabolism in the plant are among the main factors involved in stress tolerance mediated by symbiotic microbes (Vandenkoornhuyse et al. 2001; Gehring et al. 2017). In this regard, plants can recognize microbial stimuli and microbial associated molecules by certain transmembrane receptors. Recognition translates into the activation of an induced systemic resistance (ISR) boosted by the cross-talking among different hormone pathways (Van Wees et al. 2008; Morán-Diez et al. 2012). The derived molecular signals spread towards distal parts inducing a “primed” state in the plant and preparing its immune system to better combat subsequent pathogen attacks. The priming effect is characterized by a faster and stronger activation of defenses upon infection, resulting in an enhanced resistance level without a constant activation of the defense pathways (Martínez-Medina et al. 2016). Fungal endophytes are among the wide variety of microorganisms inducing a priming effect. For example, *Trichoderma* spp. and *Piriformospora indica* have shown the ability to reduce pathogen incidence in tomato and barley, respectively (Waller et al. 2008; Martínez-Medina et al. 2013; Pescador et al. 2022).

Apart from the molecular mechanisms induced in the plant by fungal endophytes, increased resistance to pathogens can be also exerted by direct inhibition of pathogen growth in plant tissues (Terhonen et al. 2019). Fungal endophytes of forest trees are receiving growing interest as biocontrol agents against a wide range of pathogens (Witzell et al. 2014; Romeralo et al. 2015; Rabiey et al. 2019). Concerning elms, DED-resistant plant material has been used to assess elm microbiome composition and disentangle differences in fungal communities between resistant and susceptible genotypes (Martín et al. 2013b). Recently, an association has been reported between the abundance of certain members of the *U. minor* core mycobiome and the degree of host resistance to DED (Macaya-Sanz et al. 2020). Certain elm endophytes have demonstrated biocontrol potential against DED (Martínez-Arias et al. 2021c), and activity in their hosts as growth stimulators and abiotic stress reducers (Martínez-Arias et al. 2021b). Furthermore, novel and low-cost screening techniques with early developed in vitro elm plants have been developed to shorten the long breeding cycles and the large experimental areas required with the current elm breeding methods (Martín et al. 2019a; Martínez-Arias et al. 2021b). By using this in vitro technique, the present work aimed to evaluate: i) the early defense response of resistant and susceptible elm genotypes to *O. novo-ulmi*, and ii) how this response is produced in elms inoculated with a core endophyte (*Cystobasidium* sp.) before exposure to *O. novo-ulmi*. It was hypothesized that a different molecular response would be detected according to the DED-resistance level of the genotype during early developmental stages. Moreover, given that *Cystobasidium* sp. was classified as plant growth-promoting yeast (Joubert and Doty 2018) and is within the group of endophytes associated with resistant elm genotypes (Macaya-Sanz et al. 2020), it was also postulated that its presence in elm tissues could trigger enhanced defense responses against *O. novo-ulmi*.

## Material and Methods

### Plant material

Four Spanish *Ulmus minor* genotypes were used for the experiment. The genotypes M-DV2.3 (Dehesa de Amaniel, Madrid) and V-AD2 (Ademúz, Valencia) were selected as representatives of resistant genotypes (i.e. < 10% of the crown showing foliar wilting 60 days after inoculation with *O. novo-ulmi*), while the genotypes M-DV1 (Dehesa de la Villa, Madrid) and VA-AP38 (Arrabal del Portillo, Valladolid) were selected for being highly susceptible (i.e. > 80% of leaf wilting) according to previous susceptibility tests performed by the Spanish elm breeding program (Martín et al. 2015). VA-AP38 belongs to the Atinian clone (*U. procera*) a highly DED-susceptible genotype spread mostly throughout Spain and England since Roman times (Gil et al. 2004).

In vitro plant production was performed as described by Martín et al. (2019a). First, buds from adult trees were sterilized with 70% ethanol for 3 min followed by 10 min incubation in 1.5% sodium hypochlorite, and then rinsed three times in distilled sterilized water. For the stimulation of aerial organs, buds were cultured in DKW basal medium (pH 5.7; Driver & Kuniyuki, 1984) gelled with 8 g l^-1^ agar and supplemented with 2.5 µM benzyl-6-amino purine. Then, aerial explants were transferred to an in vitro pot with DKW-Agar medium supplemented with 1.3 µM indole-3-butyric acid to promote differentiation of root tissue. Cultures were kept in a growth chamber at 25 °C with 16 h photoperiod using fluorescent white light. Once developed, individual plants were transferred to the experimental system and assigned to the different treatments (see below).

### Inoculum preparation

*Ophiostoma novo-ulmi* inoculum was produced by using the SOM-1 isolate, identified as *O. novo-ulmi ssp. americana* by Martín et al. (2019a). Fungal plugs were grown on malt extract agar (MEA) for 7 days. Then, mycelial fragments from the colony edge were grown in Erlenmeyer flasks with Tchernoff’s liquid medium (Tchernoff 1965) at 22 °C in the dark under constant shaking to induce sporulation. Three days later, the liquid suspension was filtered to remove germinated spores and centrifuged to collect the spores. Tchernoff medium was removed and substituted by sterile distilled water. The spore concentration was set at 4 · 10^7^ blastospores ml^-1^ using a hemocytometer.

A fungal endophyte identified as *Cystobasidium* sp. (deposited in the Spanish Type Culture Collection (CECT) under the reference CECT13192; Martínez-Arias et al., 2021b) was selected for this study. This yeast was isolated from twigs of a DED-resistant *U. minor* genotype growing in a conservation plot at “Puerta de Hierro” Forest Breeding Center (Madrid, Spain) and was coded as P5. To obtain P5 inoculum, yeast cells were refreshed twice by growing them on yeast extract agar (YEA). Then, the cells were dragged from the agar by using a sterile spatula and suspended in sterile distilled water. The final concentration was adjusted to 4 · 10^7^ cells ml^-1^ using a hemocytometer.

### Experimental design and sampling

Plants from in vitro propagation were assigned to four different treatments and transferred individually to sterile glass culture vessels. In vitro and sterile conditions were maintained during the whole experiment to avoid foreign contaminations. The experiment comprised 64 in vitro plants, i.e., 16 plants per genotype and 4 biological replicates per genotype and treatment. The four treatments were: i) control plants inoculated with sterile distilled water (C treatment), ii) plants inoculated with *O. novo-ulmi* spore suspension (Oph treatment), iii) plants inoculated with P5 endophyte cell suspension (P5 treatment) and iv) plants pre-inoculated with P5 a week before *O. novo-ulmi* inoculation (P5+Oph treatment). Inoculations were performed by cutting the root system at 3 cm from the callus and maintaining it submerged in the treatment suspension for 1 min (Martín et al. 2019a). Then, plants were transferred into individual sterile glass vessels (6 cm diameter x 9.5 cm height) containing 50 g of autoclave sterilized sand as substrate, and supplemented with 10 ml of MS nutritive medium (Murashige and Skoog 1962). Plants were grown in a chamber at 25/20 °C day/night temperatures, with a 16 h photoperiod and relative air humidity of 40%.

One week after inoculation, the four biological replicates per genotype and treatment were extracted from the culture vessel (Fig. 1). The presence of new roots was visually evaluated according to the following scale: (-) absence, (+) presence, (++) high presence. Shoots and roots were immediately detached and frozen using liquid nitrogen for being stored at −80 °C until use for molecular and biochemical analyses (see below). For all the analyses, the frozen plant material was ground to a fine powder using a Mixer mill MM 400 (Retsch GmbH, Haan, Germany) set to 30 Hz for 30 sec.

**Figure 1.**
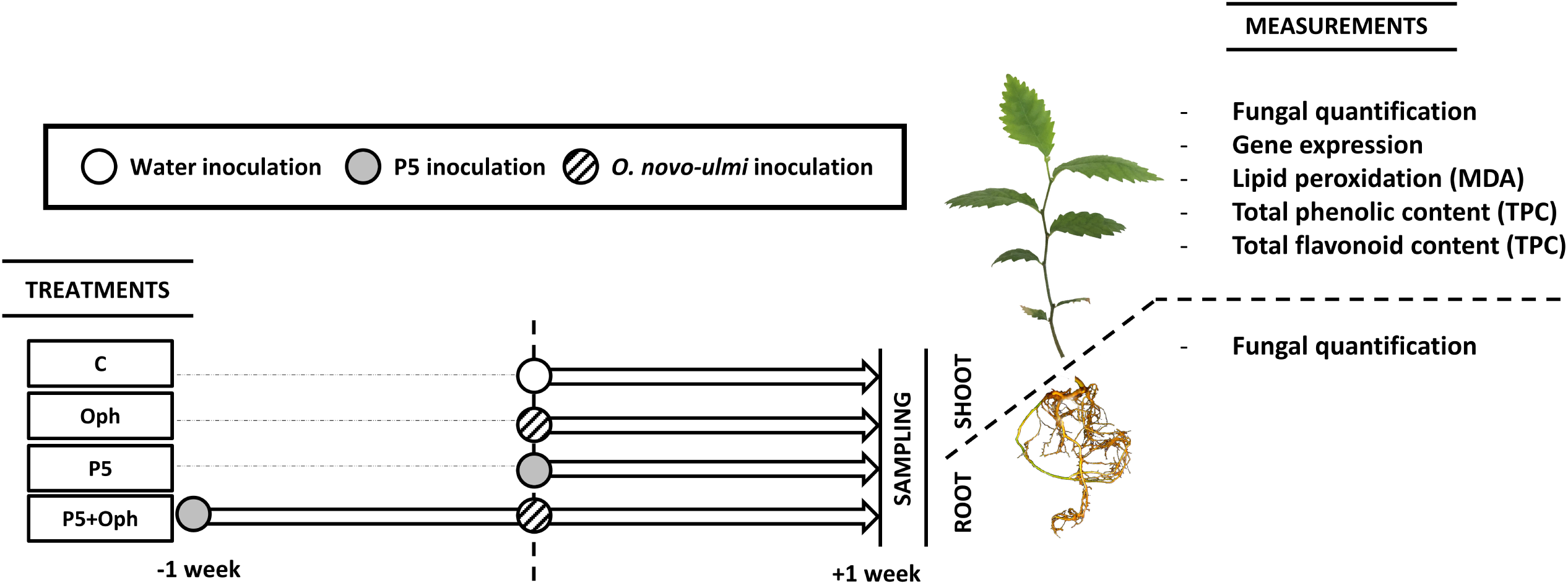
Experimental design and scheme of sampling for the in vitro-grown elm plantlets used in the study and the subsequent measurements performed in each organ. (MDA = quantification of malondialdehyde).

### In planta fungal detection

Fungal colonization ability from roots to shoot tissues was evaluated. DNA was extracted from 50 mg of the root or shoot powder material using the DNeasy plant kit (Qiagen, Hilden, Germany). Specific primer sequences were designed within the ITS2 region of *O. novo-ulmi* and P5 rRNA using Primer3 Version 0.4.0 (http://bioinfo.ut.ee/primer3-0.4.0/primer3/) (Table S1). Fungal DNA quantifications were performed with SYBR Green Master Mix (Thermo Fisher Scientific, Waltham, MA, USA) in a ViiA™ 7 Real-Time PCR system (Applied Biosystems, Foster City, CA, USA) with a standard amplification protocol. Fungal colonization was determined by the 2^-ΔCt^ method (Schmittgen and Livak 2008) by subtracting the raw threshold cycle (Ct) values of *O. novo-ulmi* or P5 ITS2 from those of *U. minor* 18S-rRNA. The amplification results were expressed as *O. novo-ulmi* and P5 presence in each sample relative to average presence of both organisms in roots of Oph- and P5-inoculated M-DV2.3 plantlets, respectively.

### Gene expression analysis

The expression of twelve defense-related genes was analyzed by quantitative reverse transcription polymerase chain reaction (qRT-PCR) in shoot tissues (Table S1). These genes were selected on the basis of the annotation results from a *U. minor* transcriptome analysis previously performed in our group (Perdiguero et al. 2015). Approximately 100 mg of shoot material was used for RNA extraction using the plant RNA isolation kit Spectrum Plant total RNA Kit (Sigma-Aldrich Chemie GmbH., Steinheim, Germany). The obtained RNA was treated with DNAse (Thermo Fisher Scientific, Waltham, MA, USA). First-strand cDNA was synthesized from 1 µg of total RNA from each sample using RevertAid H minus Reverse Transcriptase (Thermo Fisher Scientific, Waltham, MA, USA) according to the manufacturer instructions. Quantitative RT-PCRs were performed using the SYBR Green Master Mix (Thermo Fisher Scientific, Waltham, MA, USA) in a ViiA™ 7 Real-Time PCR system (Thermo Fisher Scientific, Waltham, MA, USA) with a standard amplification protocol. Three technical replicates were processed for each biological replicate. Relative quantification of specific mRNA levels was performed using the comparative method of Livak and Schmittgen (2001). Expression values were normalized using the housekeeping gene 18S-rRNA (Ref) for *U. minor*. Gene expression was considered to be up-or down-regulated if fold-change values were ≥ 1.5 or ≤ 0.7, respectively, besides being significantly different from control plants (see Statistical analysis section).

### Total phenolic and flavonoid contents

For the quantification of total phenolic and flavonoid contents (TPC and TFC, respectively), around 20 mg of powdered shoot material was extracted in 1 ml of 95% methanol under constant shaking in a Precellys Evolution mixer mill (Bertin Instruments, Montigny-le-Bretonneux, France), then incubated at room temperature for 48 h in the dark. After this time, samples were centrifuged, and supernatants were recovered. TPC was quantified using the Folin-Ciocalteu reagent (F-C) according to a microplate-adapted protocol described by Ainsworth & Gillespie (2007). On the other hand, TFC was determined by the colorimetric method described in Barreira et al. (2008), adapting volumes for a microplate reader protocol. Results were expressed as mg of gallic acid and quercetin equivalents, for TPC and TFC respectively, per gram of fresh weight of sample.

### MDA quantification

Lipid peroxidation was determined as an oxidative stress parameter by measuring the malondialdehyde (MDA) content on 50 mg of powdered shoot material. MDA concentration was determined according to Ortega-Villasante et al. (2005). Results were expressed as nmol of MDA per gram of fresh weight of sample.

### Statistical analysis

All the dependent variables, except gene expression, were analysed using two-way analysis of variance (ANOVA), with genotype and treatment and their interaction as between-subject factors, followed by Fisher’s Least Significance Difference (LSD) post-hoc tests to differentiate means when ANOVA showed significant effects (*p* < 0.05). Fold change values of gene expression analyses were compared between treatments within each genotype using one-way ANOVA, followed by Fisher’s Least Significance Difference (LSD) post-hoc test to differentiate means (*p* < 0.05 and *p* < 0.1). When needed, data were log-or arcsine-transformed prior to analysis to comply with normality and homoscedasticity assumptions. All the analyses were run using STATISTICA version 8.0 (StatSoft, Tulsa, OK, USA).

## Results

### Ophiostoma novo-ulmi *detection* in planta

Different patterns of *O. novo-ulmi* presence were observed among genotypes in root and shoot tissues (*p* < 0.01; Table 1). In the whole plant, *O. novo-ulmi* abundance was lower in plants pre-inoculated with P5 endophyte than in plants not inoculated with P5 (*p* < 0.01; Table 1). *O. novo-ulmi* colonization was more successful in roots (inoculation organ) than in shoots (Fig. 2). Focusing on Oph-treated plants, the resistant genotype M-DV2.3 showed the lowest pathogen presence in root tissues (*p* < 0.05; Fig. 2B). Conversely, in shoot tissues, the genotypes M-DV2.3 and M-DV1 showed higher *O. novo-ulmi* presence than V-AD2 and VA-AP38 (*p* < 0.05; Fig. 2A). Furthermore, pre-inoculation with P5 markedly reduced *O. novo-ulmi* presence in M-DV2.3 shoots and in VA-AP38 roots (*p* < 0.05; Figs. 2A, B).

**Table 1.** Results of factorial ANOVA of variables measured in Ulmus minor plantlets. The effect of Genotype, Treatment and their interaction, Genotype × Treatment, was studied. Red and bold numbers in p-value indicate statistically significant differences at p < 0.05. (TPC: Total Phenolic Content; TFC: Total Flavonoids Content; MDA: malondialdehyde).

| Variable | Effect | Sum of squares (SS) | Degrees of freedom (df) | Mean squares (MS) | F-ratio | *p*-value |
| --- | --- | --- | --- | --- | --- | --- |
| ***O. novo-ulmi* presence in shoots** | Genotype | 0.214 | 3 | 0.071 | 7.813 | **0.001** |
|  | Treatment | 0.024 | 1 | 0.024 | 2.608 | 0.122 |
|  | Genotype*Treatment | 0.037 | 3 | 0.012 | 1.344 | 0.288 |
| ***O. novo-ulmi* presence in roots** | Genotype | 69.87 | 3 | 23.29 | 8.966 | **0.001** |
|  | Treatment | 22.54 | 1 | 22.54 | 8.679 | **0.009** |
|  | Genotype*Treatment | 22.23 | 3 | 7.41 | 2.853 | 0.066 |
| ***O. novo-ulmi* presence in whole plant** | Genotype | 75.60 | 3 | 25.20 | 10.540 | **0.000** |
|  | Treatment | 24.45 | 1 | 24.45 | 10.225 | **0.005** |
|  | Genotype*Treatment | 23.42 | 3 | 7.81 | 3.265 | **0.046** |
| **P5 presence in shoots** | Genotype | 0.003 | 3 | 0.001 | 0.842 | 0.485 |
|  | Treatment | 0.01 | 1 | 0.01 | 7.351 | **0.012** |
|  | Genotype*Treatment | 0.009 | 3 | 0.003 | 2.296 | 0.104 |
| **P5 presence in roots** | Genotype | 16.12 | 3 | 5.373 | 7.005 | **0.002** |
|  | Treatment | 1.29 | 1 | 1.285 | 1.676 | 0.209 |
|  | Genotype*Treatment | 3.45 | 3 | 1.149 | 1.498 | 0.243 |
| **P5 presence in whole plant** | Genotype | 16.53 | 3 | 5.510 | 7.088 | **0.002** |
|  | Treatment | 1.20 | 1 | 1.195 | 1.538 | 0.228 |
|  | Genotype*Treatment | 3.04 | 3 | 1.015 | 1.305 | 0.298 |
| **TPC** | Genotype | 4.194 | 3 | 1.398 | 6.68 | **0.001** |
|  | Treatment | 8.091 | 3 | 2.697 | 12.89 | **0.000** |
|  | Genotype*Treatment | 2.033 | 9 | 0.226 | 1.08 | 0.398 |
| **TFC** | Genotype | 2.194 | 3 | 0.731 | 3.232 | **0.033** |
|  | Treatment | 4.918 | 3 | 1.639 | 7.246 | **0.001** |
|  | Genotype*Treatment | 1.075 | 9 | 0.119 | 0.528 | 0.845 |
| **MDA level** | Genotype | 10300 | 3 | 3419 | 1.484 | 0.232 |
|  | Treatment | 57200 | 3 | 19100 | 8.276 | **0.000** |
|  | Genotype*Treatment | 51000 | 9 | 5664 | 2.458 | **0.023** |

**Figure 2.**
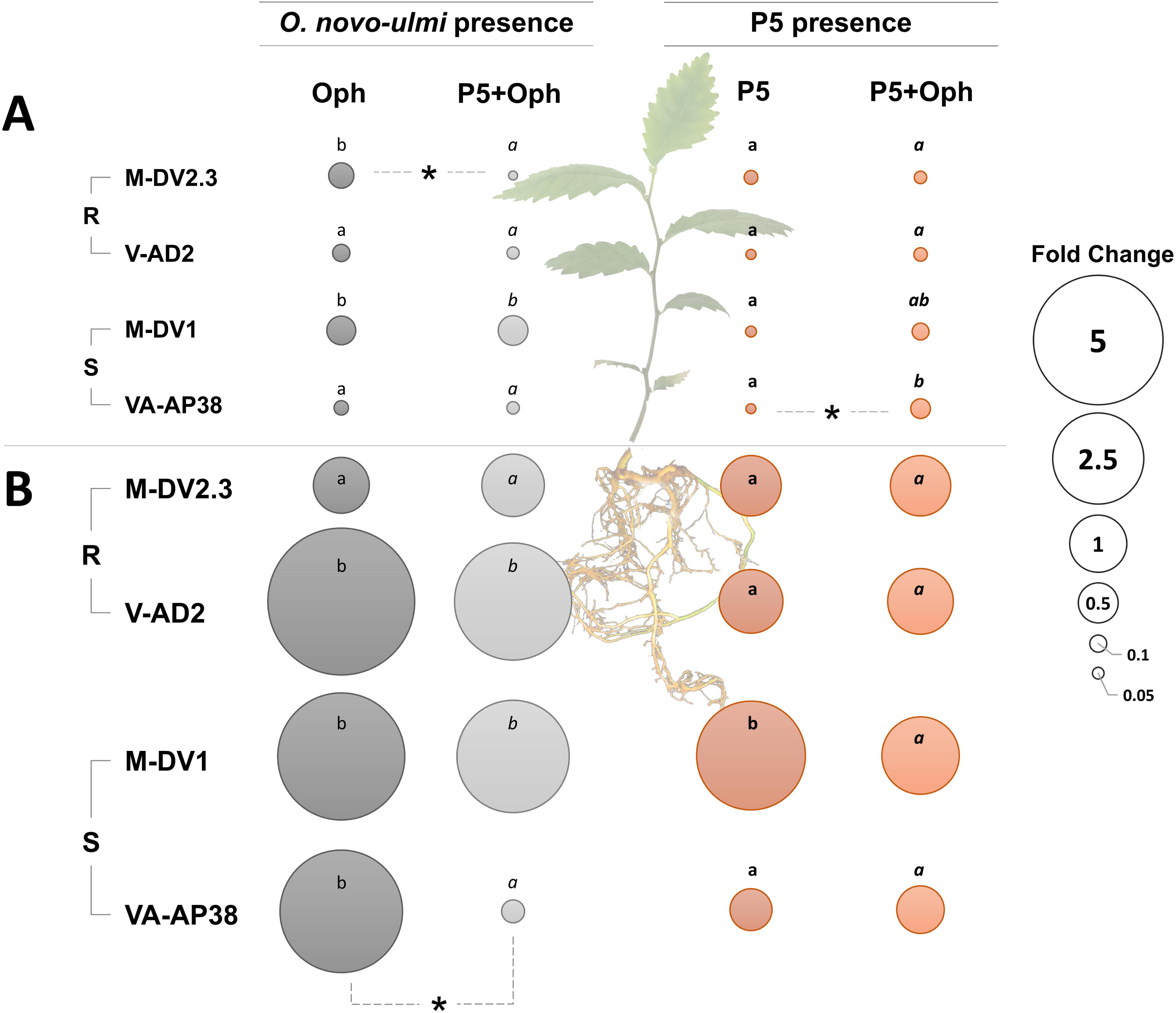
Relative presence of *Ophiostoma novo-ulmi* and P5 endophyte in shoots (**A**) and roots (**B**) of resistant (R) and susceptible (S) *Ulmus minor* genotypes. Results are shown as circles representing the average fold change of fungal presence relative to roots of Oph-or P5-treated plantlets from the resistant genotype M-DV2.3. Within each treatment, different letters indicate significant differences between genotypes, with independence for each plant organ (*p* < 0.05; Fisher’s LSD test). Differences in *O. novo-ulmi* presence (grey spheres) are represented by normal letters in Oph treatment and by italic letters in P5+Oph treatment. Regarding P5 presence (pink spheres), bold letters correspond to P5 treatment and bold italic letters correspond to P5+Oph treatment. In each genotype, significant pairwise comparisons in Oph vs P5+Oph or in P5 vs P5+Oph are indicated with asterisks (*p* < 0.05; Fisher’s LSD test).

### *Endophyte detection* in planta

The colonization of the P5 yeast was rather similar in shoots of all elm genotypes, however, in M-DV1 roots the presence of this endophyte was 3.0-fold higher than in the rest (*p* < 0.05; Fig. 2B; Table 1). When plants were inoculated with both the endophyte and the pathogen, P5 presence did not diminish or even increased in comparison with plants inoculated with P5 only (*p* < 0.05; Fig. 2A; Table 1).

### Gene expression analysis

The expression of twelve genes related to plant defense responses was evaluated in the shoot tissue. Each of the four elm genotypes evaluated showed a unique gene up-regulation pattern induced by *O. novo-ulmi* inoculation (Fig. 3). Yet, in all genotypes the pre-inoculation with P5 endophyte lowered the number of up-regulated genes after *O. novo-ulmi* inoculation. For instance, in VA-AP38 the *O. novo-ulmi* inoculation up-regulated seven genes (*p* < 0.05), but when this genotype was pre-inoculated with P5 only one gene was up-regulated (*p* < 0.05) (Fig. 3). Interestingly, the pre-inoculation with P5 increased the number of down-regulated genes after *O. novo-ulmi* inoculation in the two resistant genotypes (Fig. 3).

**Figure 3.**
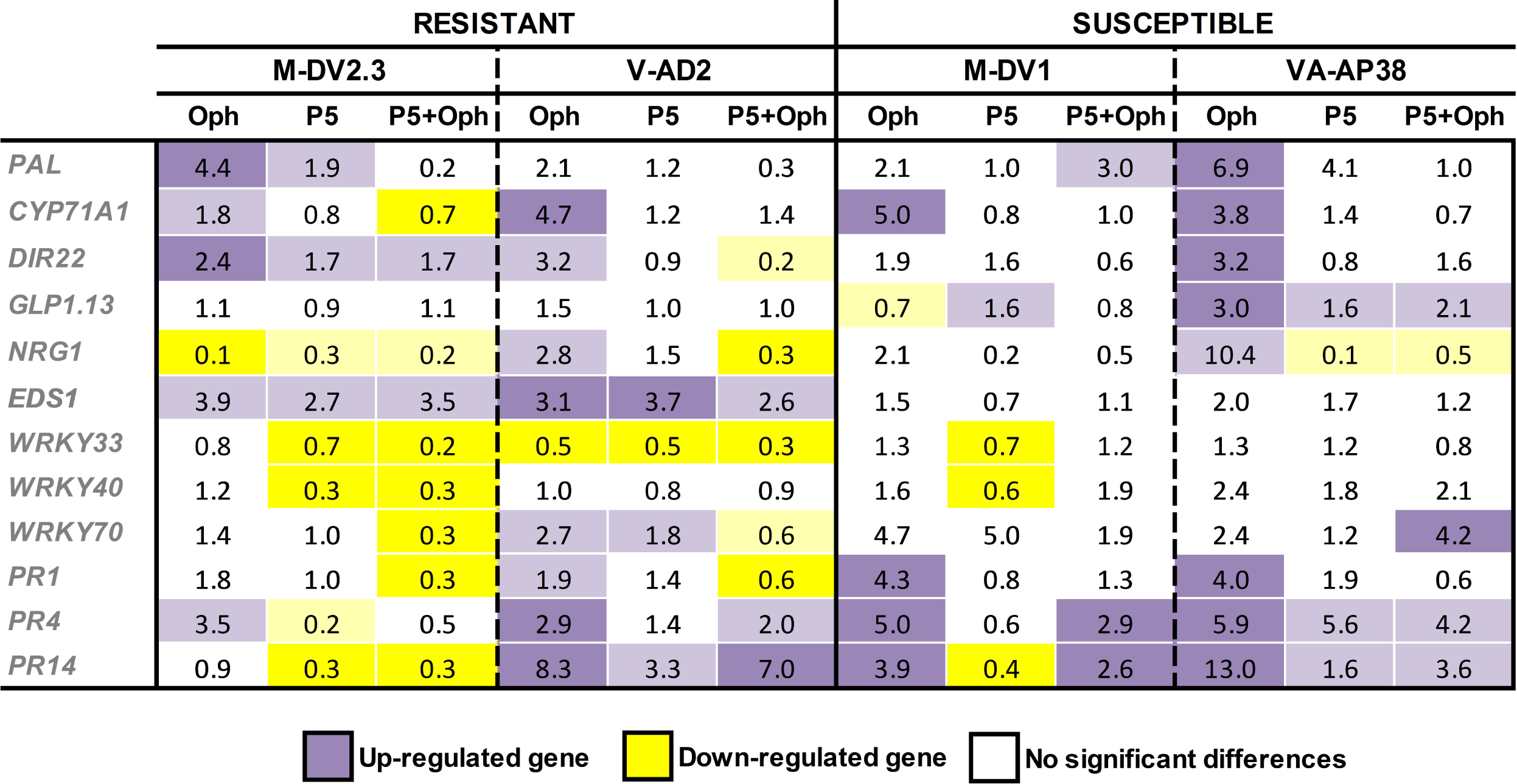
Transcriptional qRT-PCR profile of selected genes in shoots of resistant and susceptible *Ulmus minor* genotypes. Values are means of four independent biological replicates. Relative expression was normalized to the *U. minor* reference gene 18S-rRNA. Genes were considered to be significantly up-or down-regulated for a genotype if fold-change values were ≥ 1.3 (purple-colored) or ≤ 0.75 (yellow-colored) respectively, besides being statistically significant (dark colors for *p* < 0.05 and light colors for *p* < 0.1; Fisher’s LSD test).

The two DED-resistant genotypes showed an up-regulation of the *EDS1* gene after *O. novo-ulmi*, P5 endophyte and their combined inoculation, while the expression of this gene did not change in the two susceptible genotypes after the same treatments (Fig. 3).

The susceptible genotype VA-AP38 showed the highest number of up-regulated genes in response to *O. novo-ulmi* inoculation with seven out of the twelve studied genes up-regulated (58.3%; *p* < 0.05) (Fig. 3). This percentage was only 16% in M-DV2.3 and 33.3% in both V-AD2 and M-DV1 genotypes.

The sole inoculation of P5 endophyte induced different responses in each genotype, but in general, a reduced up-regulation of defense-related genes was observed when compared to the pathogen inoculation (Fig. 3).

### Biochemical analyses

The level of phenolic metabolites (flavonoids and total phenolics) was measured in shoot tissue to evaluate the plant chemical response after inoculations. Both, the elm genotype and the treatment (control, endophyte, pathogen and endophyte+pathogen inoculations) performed significant effects on total flavonoid content (TFC) and total phenolic content (TPC) (*p* < 0.05; Table 1). The *O. novo-ulmi* inoculation increased TPC levels in all genotypes, except in the resistant M-DV2.3 (Fig. 4). The endophyte inoculation alone did not alter TFC or TPC levels. However, the pre-inoculation of the endophyte attenuated the accumulation of phenolic metabolites in response to *O. novo-ulmi* inoculation (Fig. 4).

**Figure 4.**
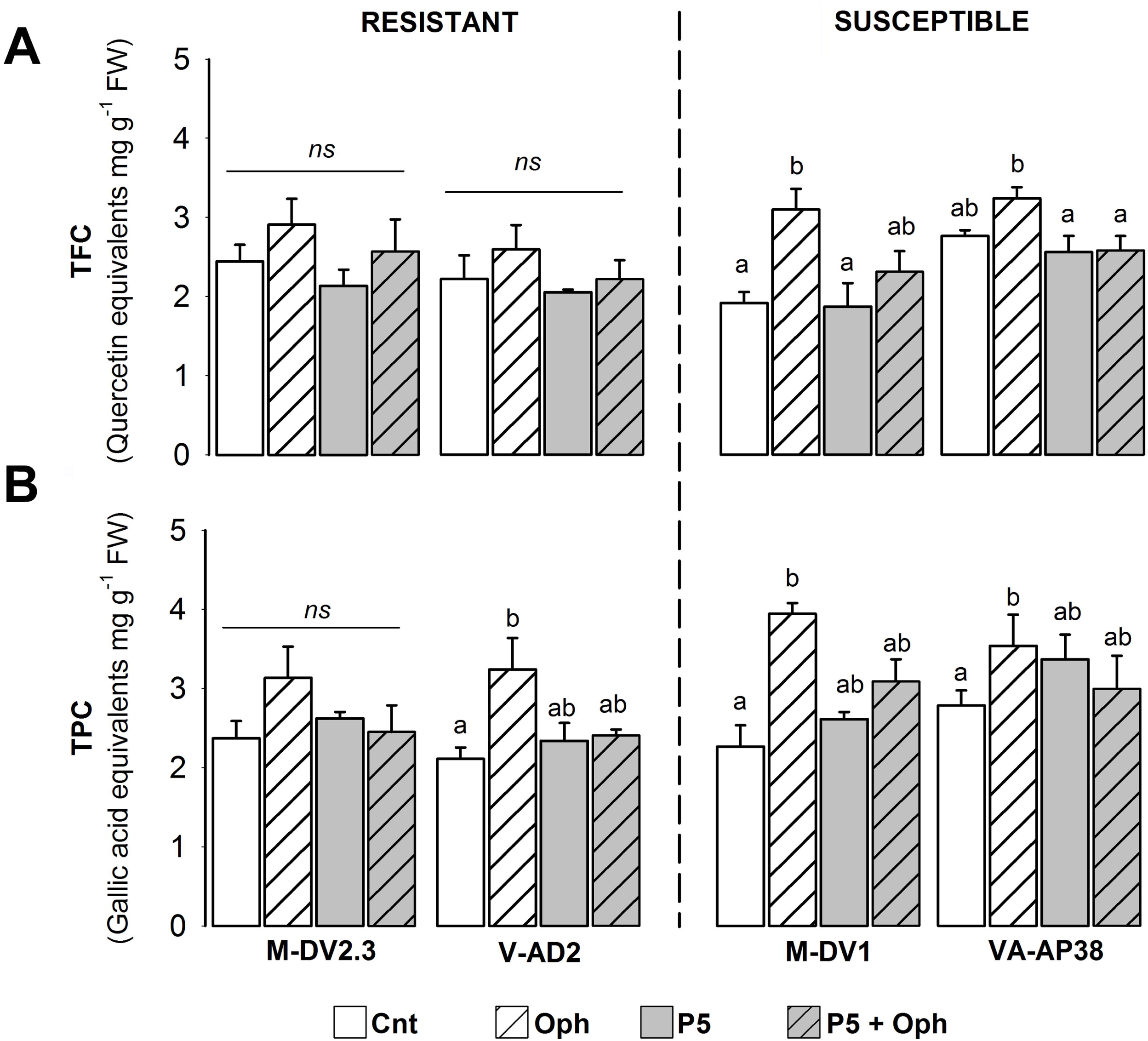
Total flavonoid content (TFC) expressed as quercetin equivalents (**A**), and total phenolic content (TPC) expressed as gallic acid equivalents (**B**), measured in shoots of elm plantlets from DED-resistant and DED-susceptible *Ulmus minor* genotypes. Within each genotype, different letters indicate significant differences between treatments (*p* < 0.05; Fisher’s LSD test; *ns* = non-significant differences).

Oxidative stress in plants after pathogen and endophyte inoculations was estimated through the level of lipid peroxidation (MDA content) in shoot tissues (Fig. 5). Lipid peroxidation in DED-resistant genotypes was not altered by pathogen or endophyte inoculation. Meanwhile, the pre-inoculation with the endophyte diminished the redox imbalance induced by the pathogen in DED-susceptible genotypes (*p* < 0.05; Fig. 5).

**Figure 5.**
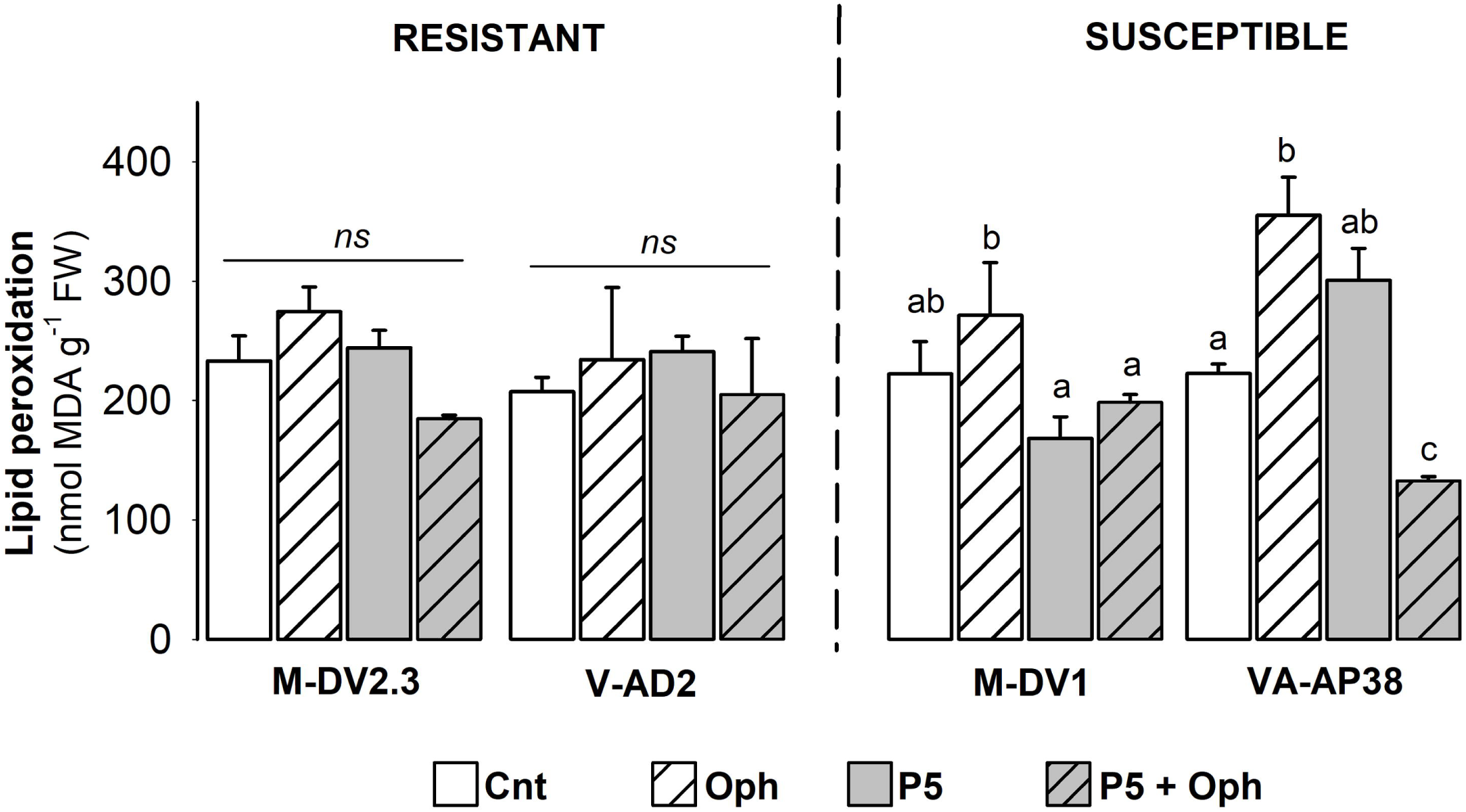
Lipid peroxidation measured through the quantification of malondialdehyde (MDA) produced in shoots of elm plantlets from DED-resistant and DED-susceptible *Ulmus minor* genotypes. Within each genotype, different letters indicate significant differences between treatments (*p* < 0.05; Fisher’s LSD test; *ns* = non-significant differences).

## Discussion

### *P5 endophytic yeast hindered colonization of elm plantlets by* O. novo-ulmi

The present work reveals intraspecific trends in *U. minor* responses to plant colonization by the DED pathogen *O. novo-ulmi* and by an endophytic yeast of the genus *Cystobasidium* (coded as P5). We found that plant colonization by the pathogen was dependent on the host genotype, with no straightforward relation with the DED resistance level of the host. One genotype stood out for its lower *O. novo-ulmi* abundance in root tissue: the DED-resistant M-DV2.3 (Fig. 2). The reduced pathogen proliferation in this genotype may be indicative of and partly explain its high DED-resistance level. This result supports a recent work where higher *O. novo-ulmi* dispersal was observed in M-DV1 than in M-DV2.3 plants (Martínez-Arias et al. 2021a), indicating a consistent behavior of M-DV2.3 in limiting *O. novo-ulmi* spread when compared to other genotypes, possibly because this genotype develops vessels of smaller diameter (Pita et al. 2018) and length (authors, unpublished results). In turn, colonization of plants by P5 was rather similar in all elm genotypes, with the exception of the susceptible M-DV1, whose roots showed higher presence of P5. Interestingly, P5 pre-inoculation reduced the abundance of *O. novo-ulmi* in the whole plant (*p* = 0.005; Table 1), while the abundance of P5 did not change after *O. novo-ulmi* inoculation (*p* = 0.228; Table 1). This result shows that the pathogen was not able to displace the endophyte from plant tissues during the experiment. The lower pathogen presence in plants already colonized by the endophyte could directly reduce the impact of the pathogen on plant physiology and metabolism. However, the effect of the endophyte-mediated lowering of pathogen presence also varied with the plant genotype (Fig. 2), revealing the complexity of host-endophyte-pathogen interactions and that the functions of endophytes on plants cannot always be generalized at the host species level.

### P5 attenuated plant defense responses to the pathogen

The pre-inoculation of the endophyte diminished the number of up-regulated plant defense-related genes induced by *O. novo-ulmi* (Fig. 3). The lower presence of pathogen cells in plant tissues colonized by the endophyte probably attenuated the intensity of plant responses against the pathogen. In addition, the pre-inoculation of the endophyte increased the number of down-regulated genes in response to the pathogen in the resistant genotypes. This endophyte-host interaction can be interpreted in different ways. On the one hand, the down-regulation of defense-related genes after *O. novo-ulmi* inoculation might indicate a mechanism to weaken the plant defense system to prevent a strong reaction against the pathogen, similarly as occurs in mutualistic interactions of microbes and plants, where SA-mediated plant defenses are inhibited or attenuated to allow these organisms to live in their tissues (Zamioudis and Pieterse 2012; Plett and Martin 2018). By down-regulating defense responses, the plant could invest resources in other processes than defense (e.g., root or shoot growth), which could promote plant tolerance to the infection (Martín et al. 2019b). This hypothesis agrees with the fact that down-regulation of defense-related gene expression after *O. novo-ulmi* inoculation was observed in the resistant genotypes only, which are able to continue growing after infection (Martín et al. 2019b). On the other hand, the down-regulation of defense-related genes induced by P5 inoculation suggests that attenuation of pathogen-induced stress may be also regulated at molecular level, particularly in the resistant genotypes. It should be acknowledged that a previous metabarcoding study with adult elm trees concluded that operational taxonomic units within the class Cystobasidiomycetes (which includes the P5 isolate) are more abundant in resistant than in susceptible elm genotypes under field conditions (Macaya-Sanz et al. 2020). Although this trend was not confirmed under the very different in vitro experimental conditions of the present work, our results suggest that resistant genotypes established a closer molecular interaction with the endophytic yeast than susceptible genotypes, leading to down-regulation of a higher number of defense-related genes after pathogen infection than susceptible trees. However, the limited number of genes explored in this work makes it necessary to confirm this trend by analyzing a larger array of genes, e.g., through high throughput sequencing techniques (e.g., RNAseq).

### P5 mitigated the pathogen-induced stress in DED-susceptible clones

Besides the observed host-endophyte-pathogen interaction in defense gene regulation, other plausible mechanisms by which P5 endophyte attenuated the stress caused by *O. novo-ulmi* in elm plants are: i) ability to control the levels of oxidative stress induced by the pathogen, ii) promotion of root growth; and iii) direct antagonism and/or niche competition against *O. novo-ulmi*. Regarding the first mechanism, P5 pre-inoculation lowered the levels of lipid peroxidation (MDA content) in susceptible trees after pathogen infection (Fig. 5). The increase of oxidative stress and phenolic metabolites after *O. novo-ulmi* infection was mainly observed in susceptible genotypes, similarly as observed in a previous work (Martín et al. 2019a). Total phenolics and flavonoids accumulation induced by *O. novo-ulmi* can be related to an attempt of the plant to limit pathogen spread (Ouellette and Rioux 1992; Witzell and Martín 2008) and to counterbalance the oxidative damage derived from the incompatible interaction between the plant and the pathogen (Shalaby and Horwitz 2015). In this sense, no increase of oxidative stress was observed in response to P5 inoculation, which evidences the non-pathogenic nature of this endophyte towards elm. Concerning the second mechanism (promotion of root growth), a positive effect of P5 on root growth stimulation was observed in P5-inoculated plants (Figure S1). Higher formation of fine roots may help the plant sustaining water uptake and hydraulic functioning during pathogen invasion of xylem conduits. The production of the auxin indole acetic acid (IAA) by P5 endophyte was demonstrated in a previous work (Martínez-Arias et al. 2021c) and may contribute to the stimulation of root formation in host plants (Harman 2011; Sukumar et al. 2013). In other work, P5-inoculated elm plantlets showed higher root development accompanied with higher survival rates against abiotic stress than non-inoculated plants (Martínez-Arias et al. 2021b), evidencing the role of this yeast in counterbalancing plant stressful situations. Although the third mechanism (direct antagonism and/or niche competition against *O. novo-ulmi*) was not directly evaluated in the present work, previous analyses demonstrated that liquid filtrates from P5 reduced *O. novo-ulmi* growth, and that P5 partly overlapped with *O. novo-ulmi* in the metabolization of different carbon sources (Martínez-Arias et al. 2021c). Particularly, P5 was able to grow in presence of defensive molecules (flavonoids and other phenolic compounds) produced either by the host or the pathogen, possibly helping the yeast to compete with the pathogen within plant tissues. Furthermore, other plant-growth promoting yeasts similar to P5 have been also described as beneficial symbionts, not only by promoting plant growth but also by acting as inhibitors of phytopathogens (El-Tarabily 2004; Ignatova et al. 2015), enhancing plant defense, or performing a direct antagonism to pathogens (Calvente et al. 2001; Kalogiannis et al. 2006; Akhtyamova and Sattarova 2012; Gava et al. 2018).

### Different molecular responses evidence the multifactorial nature of DED resistance

Our results further confirm the complexity and heterogeneity of elm defense mechanisms against DED among different genotypes. In response to *O. novo-ulmi* inoculation, the two DED-susceptible genotypes and the resistant V-AD2 shared a rather similar pattern of gene regulation while the resistant M-DV2.3 showed a distinct response (Fig. 3). The different response of the two resistant genotypes to DED was evidenced, for example, by the down-regulation of *NRG1* in M-DV2.3, and the up-regulation of the same gene in V-AD2. The multigenic nature of elm resistance to DED have already been proposed in previous works (Townsend and Santamour 1993; Martín et al. 2019b) and could have important implications for elm breeding. Thus, by crossing genotypes with different and ideally complementary defense mechanisms it might be possible to enhance disease resistance in the offspring. The genes *PR4* and *CYP71A1* were found up-regulated in the four elm genotypes after pathogen infection, in agreement with previous works describing elm responses to *O. novo-ulmi* (Aoun et al. 2010; Sherif et al. 2016; Perdiguero et al. 2018). Besides these two genes, *PR1* and *PR14* were also induced by the pathogen in both DED-susceptible genotypes, being those four genes, the only ones activated in the susceptible M-DV1 among the studied genes. Yet, the other susceptible genotype (VA-AP38) up-regulated eight (i.e., 66.6%) of the analyzed genes. This genotype (representative of the so-called English elm) showed the highest expression values of all genotypes in response to pathogen inoculation. In a previous study using field-grown VA-AP38 trees, *O. novo-ulmi* inoculation stimulated the up-regulation of a large number of genes related to several metabolic pathways and biological processes, leading a tradeoff between expression of growth and defense genes (Perdiguero et al. 2018). Despite the huge transcriptional regulation, those trees were not able to stop disease progression and died. The high activation of defense pathways in this clone is clearly ineffective against the pathogen, causing a concomitant high oxidative stress. Finally, the up-regulation of *EDS1* gene after pathogen, endophyte and their combined inoculation in resistant trees but not in susceptible ones deserves further research to clarify its possible involvement in DED resistance.

## Conclusions

Numerous screening trials to find DED-resistant elm material have been performed during the decades of the Spanish Elm Breeding Program activity. As the evaluation of adult trees is very space- and time-consuming, the development of early detection tools in which these limiting factors disappear is a key step to progress in elm restoration. The potential of early screening methods using *Ulmus minor* in vitro plantlets has been previously reported (Martín et al. 2019a), showing some distinctive traits associated with DED resistance. Biochemical parameters, such as MDA and chlorophyll contents, and biometric parameters such as shoot growth were useful to distinguish between resistant and susceptible elm plantlets. Our study provides further evidence of the usefulness of the in vitro system for the early detection of DED-resistant genotypes, but also to investigate plant-endophyte symbioses. The MDA level was confirmed as a key parameter associated with DED-susceptibility. Similarly, an accumulation of plant phenolics in response to *O. novo-ulmi* was identified as a general response to the pathogen, especially in susceptible trees. The pre-inoculation of the P5 endophyte displayed ameliorative effects against *O. novo-ulmi*, evidenced by lower pathogen abundance, reduced up-regulation of plant defenses, and lower levels of oxidative stress. Root growth promotion induced by P5 was also possibly important in the maintenance of water uptake during DED infection.

## Supplementary Data

**Table S1**. List of genes analysed by quantitative-PCR.

**Figure S1**. Presence of new roots in *Ulmus minor* plantlets one week after inoculation.

## Acknowledgments

We highly thank Prof. Corné M.J. Pieterse of Plant-Microbe Interactions group at Utrecht University (UU; the Netherlands) for his advice and the use of his lab facilities. We also thank David Medel Cuesta for his technical assistance with in vitro elm propagation.

## References

Ainsworth EA, Gillespie KM (2007) Estimation of total phenolic content and other oxidation substrates in plant tissues using Folin–Ciocalteu reagent. Nat Protoc 2: 875–877

Akhtyamova N, Sattarova RK (2012) Endophytic Yeast Rhodotorula rubra Strain TG-1: Antagonistic and Plant Protection Activities. Biochem Physiol Open Access 2013 21 2: 1–6

Aoun M, Jacobi V, Boyle B, Bernier L (2010) Identification and monitoring of Ulmus americana transcripts during in vitro interactions with the Dutch elm disease pathogen Ophiostoma novo-ulmi. Physiol Mol Plant Pathol 74: 254–266

Bari R, Jones JDG (2009) Role of plant hormones in plant defence responses. Plant Mol Biol 69: 473–488

Barreira J, Ferreira I, Oliveira M, Pereira J (2008) Antioxidant activities of the extracts from chestnut flower, leaf, skins and fruit. Food Chem 107: 1106–1113

Boonekamp PM, Pieterse CMJ, Govers F, Cornelissen BJC (2019) Johanna Westerdijk (1881–1961) – the impact of the grand lady of phytopathology in the Netherlands from 1917 to 2017. Eur J Plant Pathol 154: 11–16

Brasier CM (1991) Ophiostoma novo-ulmi sp. nov., causative agent of current Dutch elm disease pandemics. Mycopathologia 115: 151–161

Calvente V, de Orellano ME, Sansone G, Benuzzi D, Sanz de Tosetti MI (2001) Effect of nitrogen source and pH on siderophore production by Rhodotorula strains and their application to biocontrol of phytopathogenic moulds. J Ind Microbiol Biotechnol 26: 226–229

Driver J, Kuniyuki A (1984) In vitro propagation of Paradox walnut rootstock. HortScience 19: 507–509

Duchesne LC, Hubbes M, Jeng RS (1986) Mansonone E and F accumulation in Ulmus pumila resistant to Dutch elm disease. Can J For Res 16: 410–412

El-Tarabily KA (2004) Suppression of Rhizoctonia solani diseases of sugar beet by antagonistic and plant growth-promoting yeasts. J Appl Microbiol 96: 69–75

Fu ZQ, Dong X (2013) Systemic Acquired Resistance: Turning Local Infection into Global Defense. Annu Rev Plant Biol 64: 839–863

Gava CAT, de Castro APC, Pereira CA, Fernandes-Júnior PI (2018) Isolation of fruit colonizer yeasts and screening against mango decay caused by multiple pathogens. Biol Control 117: 137–146

Gehring CA, Sthultz CM, Flores-Rentería L, Whipple A V., Whitham TG (2017) Tree genetics defines fungal partner communities that may confer drought tolerance. Proc Natl Acad Sci 114: 11169–11174

Gil L, Fuentes-Utrilla P, Soto Á, Cervera MT, Collada C (2004) English elm is a 2,000-year-old Roman clone. Nature 431: 1053–1053

Harman GE (2011) Multifunctional fungal plant symbionts: new tools to enhance plant growth and productivity. New Phytol 189: 647–649

Ignatova L V., Brazhnikova Y V., Berzhanova RZ, Mukasheva TD (2015) Plant growth-promoting and antifungal activity of yeasts from dark chestnut soil. Microbiol Res 175: 78–83

Joubert PM, Doty SL (2018) Endophytic Yeasts: Biology, Ecology and Applications. In: Pirttilä A, Frank A (eds) Endophytes of Forest Trees. Springer, Cham, pp. 3–14

Kalogiannis S, Tjamos SE, Stergiou A, Antoniou PP, Ziogas BN, Tjamos EC (2006) Selection and evaluation of phyllosphere yeasts as biocontrol agents against grey mould of tomato. Eur J Plant Pathol 116: 69–76

Li M, López R, Venturas M, Martín JA, Domínguez J, Gordaliza GG, Gil L, Rodríguez-Calcerrada J (2016) Physiological and biochemical differences among Ulmus minor genotypes showing a gradient of resistance to Dutch elm disease. For Pathol 46: 215–228

Liu H, Brettell LE, Qiu Z, Singh BK (2020) Microbiome-Mediated Stress Resistance in Plants. Trends Plant Sci 25: 733–743

Livak KJ, Schmittgen TD (2001) Analysis of Relative Gene Expression Data Using Real-Time Quantitative PCR and the 2−ΔΔCT Method. Methods 25: 402–408

Macaya-Sanz D, Witzell J, Collada C, Gil L, Martin JA (2020) Structure of core fungal endobiome in Ulmus minor : patterns within the tree and across genotypes differing in tolerance to Dutch elm disease. bioRxiv 2020.06.23.166454

Martín JA, Sobrino-Plata J, Coira B, Medel D, Collada C, Gil L (2019a) Growth resilience and oxidative burst control as tolerance factors to Ophiostoma novo-ulmi in Ulmus minor. Tree Physiol 39: 1512–1524

Martín JA, Sobrino-Plata J, Rodríguez-Calcerrada J, Collada C, Gil L (2019b) Breeding and scientific advances in the fight against Dutch elm disease: Will they allow the use of elms in forest restoration? New For 50: 183–215

Martín JA, Solla A, Coimbra MA, Gil L (2008) Metabolic fingerprinting allows discrimination between Ulmus pumila and U. minor, and between U. minor clones of different susceptibility to Dutch elm disease. For Pathol 38: 244–256

Martín JA, Solla A, García-Vallejo MC, Gil L (2012) Chemical changes in Ulmus minor xylem tissue after salicylic acid or carvacrol treatments are associated with enhanced resistance to Ophiostoma novo-ulmi. Phytochemistry 83: 104–109

Martín JA, Solla A, Oszako T, Gil L (2020) Characterizing offspring of Dutch elm disease-resistant trees (Ulmus minor Mill.). For An Int J For Res 1–12

Martín JA, Solla A, Ruiz-Villar M, Gil L (2013a) Vessel length and conductivity of Ulmus branches: ontogenetic changes and relation to resistance to Dutch elm disease. Trees 27: 1239–1248

Martín JA, Solla A, Venturas M, Collada C, Domínguez J, Miranda E, Fuentes P, Burón M, Iglesias S, Gil L (2015) Seven Ulmus minor clones tolerant to Ophiostoma novo-ulmi registered as forest reproductive material in Spain. iForest - Biogeosciences For 8: 172–180

Martin JA, Solla A, Woodward S, Gil L (2005) Fourier transform-infrared spectroscopy as a new method for evaluating host resistance in the Dutch elm disease complex. Tree Physiol 25: 1331–1338

Martín JA, Witzell J, Blumenstein K, Rozpedowska E, Helander M, Sieber TN, Gil L (2013b) Resistance to Dutch Elm Disease Reduces Presence of Xylem Endophytic Fungi in Elms (Ulmus spp.). PLoS One 8: e56987

Martínez-Arias C, Sobrino-Plata J, Gil L, Rodríguez-Calcerrada J, Martín JA (2021a) Priming of Plant Defenses against Ophiostoma novo-ulmi by Elm (Ulmus minor Mill.) Fungal Endophytes. J Fungi 7: 687

Martínez-Arias C, Sobrino-Plata J, Medel D, Gil L, Martín JA, Rodríguez-Calcerrada J (2021b) Stem endophytes increase root development, photosynthesis, and survival of elm plantlets (Ulmus minor Mill.). J Plant Physiol 261: 153420

Martínez-Arias C, Sobrino-Plata J, Ormeño-Moncalvillo S, Gil L, Rodríguez-Calcerrada J, Martín JA (2021c) Endophyte inoculation enhances Ulmus minor resistance to Dutch elm disease. Fungal Ecol 50: 101024

Martínez-Medina A, Fernández I, Sánchez-Guzmán MJ, Jung SC, Pascual JA, Pozo MJ (2013) Deciphering the hormonal signalling network behind the systemic resistance induced by Trichoderma harzianum in tomato. Front Plant Sci 4:

Martínez-Medina A, Flors V, Heil M, Mauch-Mani B, Pieterse CMJ, Pozo MJ, Ton J, van Dam NM, Conrath U (2016) Recognizing Plant Defense Priming. Trends Plant Sci 21: 818–822

Morán-Diez E, Rubio B, Domínguez S, Hermosa R, Monte E, Nicolás C (2012) Transcriptomic response of Arabidopsis thaliana after 24h incubation with the biocontrol fungus Trichoderma harzianum. J Plant Physiol 169: 614–620

Murashige T, Skoog F (1962) A Revised Medium for Rapid Growth and Bio Assays with Tobacco Tissue Cultures. Physiol Plant 15: 473–497

Ortega-Villasante C, Rellán-Álvarez R, Del Campo FF, Carpena-Ruiz RO, Hernández LE (2005) Cellular damage induced by cadmium and mercury in Medicago sativa. J Exp Bot 56: 2239–2251

Ouellette GB, Rioux D (1992) Anatomical and Physiological Aspects of Resistance to Dutch Elm Disease. In: Blanchette RA, Biggs AR (eds) Defense Mechanisms of Woody Plants Against Fungi. Springer, Berlin, Heidelberg, pp. 257–307

Perdiguero P, Sobrino-Plata J, Venturas M, Martín JA, Gil L, Collada C (2018) Gene expression trade-offs between defence and growth in English elm induced by Ophiostoma novo-ulmi. Plant Cell Environ 41: 198–214

Perdiguero P, Venturas M, Cervera MT, Gil L, Collada C (2015) Massive sequencing of Ulmus minor’s transcriptome provides new molecular tools for a genus under the constant threat of Dutch elm disease. Front Plant Sci 6: 1–12

Pescador L, Fernandez I, Pozo MJ, Romero-Puertas MC, Pieterse CMJ, Martínez-Medina A (2022) Nitric oxide signalling in roots is required for MYB72-dependent systemic resistance induced by Trichoderma volatile compounds in Arabidopsis. J Exp Bot 73: 584–595

Pieterse CMJ, Leon-Reyes A, Van Der Ent S, Van Wees SCM (2009) Networking by small-molecule hormones in plant immunity. Nat Chem Biol 5: 308–316

Pita P, Rodríguez-Calcerrada J, Medel D, Gil L (2018) Further insights into the components of resistance to Ophiostoma novo-ulmi in Ulmus minor: hydraulic conductance, stomatal sensitivity and bark dehydration. Tree Physiol 38: 252–262

Plett JM, Martin FM (2018) Know your enemy, embrace your friend: using omics to understand how plants respond differently to pathogenic and mutualistic microorganisms. Plant J 93: 729–746

Rabiey M, Hailey LE, Roy SR, Grenz K, Al-Zadjali MAS, Barrett GA, Jackson RW (2019) Endophytes vs tree pathogens and pests: can they be used as biological control agents to improve tree health? Eur J Plant Pathol 155: 711–729

Romeralo C, Santamaría O, Pando V, Diez JJ (2015) Fungal endophytes reduce necrosis length produced by Gremmeniella abietina in Pinus halepensis seedlings. Biol Control 80: 30–39

Schmittgen TD, Livak KJ (2008) Analyzing real-time PCR data by the comparative CT method. Nat Protoc 3: 1101–1108

Schwarz MB (1922) Das Zweigsterben der Ulmen, Trauerweiden und Pfirsichbäume: eine vergleichende-pathologische Studie. Utrecht : Oosthoek, Utrecht

Shalaby S, Horwitz BA (2015) Plant phenolic compounds and oxidative stress: integrated signals in fungal–plant interactions. Curr Genet 61: 347–357

Sherif SM, Erland LA, Shukla MR, Saxena PK (2017) Bark and wood tissues of American elm exhibit distinct responses to Dutch elm disease. Sci Rep 7: 7114

Sherif SM, Shukla MR, Murch SJ, Bernier L, Saxena PK (2016) Simultaneous induction of jasmonic acid and disease-responsive genes signifies tolerance of American elm to Dutch elm disease. Sci Rep 6: 21934

Solla A, Gil L (2002) Xylem vessel diameter as a factor in resistance of Ulmus minor to Ophiostoma novo-ulmi. For Pathol 32: 123–134

Sukumar P, Legué V, Vayssières A, Martin F, Tuskan GA, Kalluri UC (2013) Involvement of auxin pathways in modulating root architecture during beneficial plant-microorganism interactions. Plant Cell Environ 36: 909–919

Tchernoff V (1965) Methods for screening and for the rapid selection of elms for resistance to Dutch elm disease. Acta Bot Neerl 14: 409–452

Terhonen E, Blumenstein K, Kovalchuk A, Asiegbu FO (2019) Forest Tree Microbiomes and Associated Fungal Endophytes: Functional Roles and Impact on Forest Health. Forests 10: 42

Townsend AM, Santamour FS (1993) Progress in the Development of Disease-Resistant Elms. In: Sticklen M.B., Sherald J.L. (eds) Dutch Elm Disease Research. Springer New York, New York, NY, pp. 46–50

Vandenkoornhuyse P, Leyval C, Bonnin I (2001) High genetic diversity in arbuscular mycorrhizal fungi: evidence for recombination events. Heredity (Edinb) 87: 243–253

Waller F, Mukherjee K, Deshmukh SD, Achatz B, Sharma M, Schäfer P, Kogel K-H (2008) Systemic and local modulation of plant responses by Piriformospora indica and related Sebacinales species. J Plant Physiol 165: 60–70

Van Wees SC, Van der Ent S, Pieterse CM (2008) Plant immune responses triggered by beneficial microbes. Curr Opin Plant Biol 11: 443–448

Witzell J, Martín JA (2008) Phenolic metabolites in the resistance of northern forest trees to pathogens - Past experiences and future prospects. Can J For Res 38: 2711–2727

Witzell J, Martín JA, Blumenstein K (2014) Ecological Aspects of Endophyte-Based Biocontrol of Forest Diseases. In: Verma V., Gange A. (eds) Advances in Endophytic Research. Springer India, New Delhi, xpp. 321–333

Zamioudis C, Pieterse CMJ (2012) Modulation of Host Immunity by Beneficial Microbes. Mol Plant-Microbe Interact 25: 139–150

