## Supplementary material for "No priming, just fighting – endophytic yeast attenuates the defense response and the stress induced by Dutch elm disease in *Ulmus minor*": Sup. Material

**Table S1**. List of genes analysed by quantitative-PCR. Isotig numbers from *Ulmus minor* transcriptome (Perdiguero *et al.* 2015), a brief description of their annotation, primer pair sequences and length of the amplified fragments (bp) are shown.

| ID | Isotig (*Ulmus minor*) | Seq. Description | Primers pairs (5'-3') | Fragment lgth. (bp) |
|---|---|---|---|---|
| ITS2Ophio | | Internal Transcribed Spacer 2 (*O. novo-ulmi*) | CAGTACCGAACGCAAGTTCTCTCTC<br>ATGCTTAAGTTCAGCGGGTAATCCT | 150 |
| ITS2P5 | | Internal Transcribed Spacer 2 (P5 endophyte) | CAACGGATCTCTTGGCTCTC<br>AACAGACATACTCTTCGGAATACC | 136 |
| Ri18S_RT | | 18S-rRNA (*U. minor* Reference gene) | ATATGTCAAAACGACTCTCGGCAAC<br>AACTTGCGTTCAAAGACTCGATGGT | 110 |
| *CYP71A1* | isotig08772 | cytochrome p450 71a1-like | TTTCTTTCCCACAACGCTTC<br>TTTGCTGCAGGAACTGACAC | 120 |
| *DIR22* | isotig10677 | dirigent protein 22-like | GGAACACTACTTGAGACTAA<br>AGAATGAGGTTGAGAAGAG | 81 |
| *EDS1* | isotig02962 | enhanced disease susceptibility 1 | GGATTATTCTCGGGCTGTGA<br>TGCATTTTTCCAACAACCAA | 104 |
| *GLP1.13* | isotig14915 | germin-like protein subfamily 1 member 13 | GGAGTTGTTGTGGCTATT<br>ATAATCAATCCATCATTCTTCAAT | 94 |
| *NRG1* | isotig13421 | RPW8 -CNL, NRG1 (N requirement gene 1) | AAGACATTGGCAAGTTAT<br>CTCCTCATCACATATCAC | 122 |
| *PAL* | isotig14143 | phenylalanine ammonia lyase | GGTGAGGATATAGAGAAGGT<br>AACAGAGCCATTCCAATC | 93 |
| *PR1* | isotig16547 | pathogen-related protein | GAATGGCCATCTTTGAGACG<br>TCCAGTTTTGGACCCTTCAG | 106 |
| *PR4* | isotig04787 | wound-induced protein win2 | AGCCTTGATTATAGCCATT<br>ACGGTGAGAATTGTTGAT | 100 |
| *PR14* | isotig10737 | non-specific lipid-transfer protein 1-like | CTTTAATGGCGGTGTTGTCC<br>TGCCTCCGAAAACTCCATAG | 117 |
| *WRK33* | isotig11160 | probable wrky transcription factor 33-like | TCAGTGGCGTTTTCAGCATA<br>CCACCAACCACAACTCACTG | 103 |
| *WRK40* | isotig20762 | probable wrky transcription factor 40-like | TTTCTTCAACGGGAACTTGG<br>CCGTTGAAATCTTGGCCTTA | 89 |
| *WRK70* | isotig03773 | probable wrky transcription factor 70-like | TCAGACGACACATCACTT<br>CGCAGCAGAATCAGAATA | 99 |

**A**

| | RESISTANT | | SUSCEPTIBLE | |
|---|---|---|---|---|
| | M-DV2.3 | V-AD2 | M-DV1 | VA-AP38 |
| **Cnt** | + | + | + | - |
| **Oph** | - | - | - | - |
| **P5** | ++ | ++ | ++ | ++ |
| **P5+Oph** | + | - | + | + |

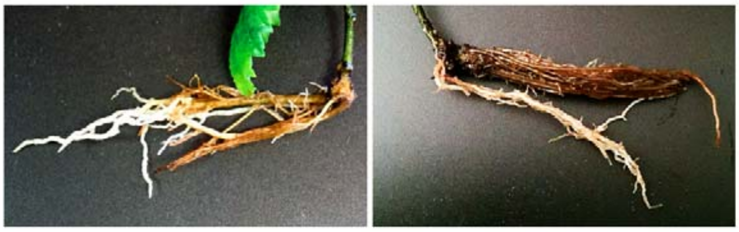

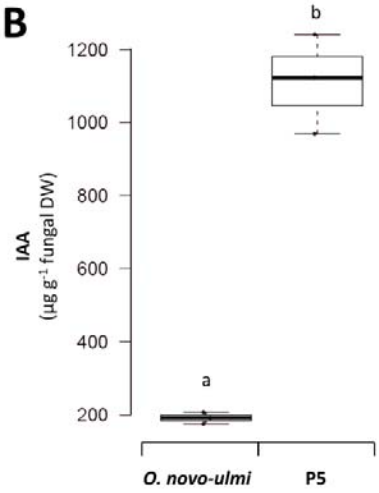


**Figure S1.** Presence of new roots in *Ulmus minor* plantlets one week after inoculation with (i) sterile water (Cnt), (ii) *Ophiostoma novo-ulmi* (Oph), (iii) P5 endophyte (P5), and (iv) P5 and *O. novo-ulmi* (P5+Oph). **A**) In the upper part, a table with visual assessment of new root growth in plantlets of each genotype. A plus sign (+) indicates presence of new roots; two plus signs (++) indicate higher amount with respect to control plants. In the lower part, two pictures of roots from M-DV2.3 (left) and M-DV1 (right) show the presence of new roots after P5 treatment. **B**) Indole-3-acetic acid (IAA) *in vitro* production by *O. novo-ulmi* and P5 endophyte after 15 days of incubation in malt extract broth containing 0.2% L-tryptophan (adapted from Martínez-Arias *et al*. 2021c).
